# The effects of prior exposure to prism lenses on *de novo* motor skill learning

**DOI:** 10.1101/2023.05.08.539850

**Authors:** Annmarie M. Lang-Hodge, Dylan F. Cooke, Daniel S. Marigold

## Abstract

Motor learning involves plasticity in a network of brain areas across the cortex and cerebellum. Such traces of learning have the potential to affect subsequent learning of other tasks. In some cases, prior learning can interfere with subsequent learning, but it may be possible to potentiate learning of one task with a prior task if they are sufficiently different. Because prism adaptation involves extensive neuroplasticity, we reasoned that the elevated excitability of neurons could increase their readiness to undergo structural changes, and in turn, create an optimal state for learning a subsequent task. We tested this idea, selecting two different forms of learning tasks, asking whether exposure to a sensorimotor adaptation task can improve subsequent *de novo* motor skill learning. Participants first learned a new visuomotor mapping induced by prism glasses in which prism strength varied trial-to-trial. Immediately after and the next day, we tested participants on a mirror tracing task, a form of *de novo* skill learning. Prism-trained and control participants both learned the mirror tracing task, with similar reductions in error and increases in distance traced. Both groups also showed evidence of offline performance gains between the end of day 1 and the start of day 2. However, we did not detect differences between groups. Overall, our results do not support the idea that prism adaptation learning can potentiate subsequent *de novo* learning. We discuss factors that may have contributed to this result.

## INTRODUCTION

Motor learning is an experience-dependent improvement in motor behavior and occurs throughout life. Uncovering methods to enhance this learning, whether through speeding up initial performance improvements, increasing overall performance, or extending the retention of a specific performance level, is important. This is because some form of motor learning is necessary for recovering function or compensation following neurological and musculoskeletal injury, to excel in athletics, and for vocational tasks.

One form of motor learning, sensorimotor adaptation, is studied with visuomotor perturbation paradigms like prism adaptation or visuomotor rotation. This learning involves a rapid, explicit, aiming-strategy component (McDougle et al. 2016) and a slower, implicit, sensory-prediction-error-driven mechanism (Maeda et al. 2017a; Tseng et al. 2007). Adaptation is characterized by the presence of negative aftereffects when the perturbation is removed or the participant attempts a different motor task, which likely reflect updating of an internal model (Bakkum et al. 2021; Bakkum and Marigold 2022; Maeda et al. 2017b; Petitet et al. 2018; Redding et al. 2005). A motor memory of this adaptation can become resistant to opposite perturbations (Bakkum and Marigold 2022; Krakauer et al. 2005; Maeda et al. 2017b) and can last for extended periods of time (>1 year) (Landi et al. 2011; Maeda et al. 2018). Prism adaptation causes activation of several regions within the posterior parietal cortex and cerebellum (Panico et al. 2020), alters post-adaptation resting-state functional connectivity in frontoparietal and cerebellar-parieto-parahippocampal networks (Schintu et al. 2020), and is improved and better retained when paired with transcranial direct current stimulation over the primary motor cortex, or simultaneously over the posterior parietal cortex and cerebellum (O’Shea et al. 2017; Panico et al. 2022). Taken together, prism adaptation involves extensive neuroplasticity.

*De novo* skill learning represents another form of motor learning. Rather than adapting an existing control policy as with prism adaptation, *de novo* learning involves creating a new control policy (Krakauer et al. 2019; Sternad 2018; Telgen et al. 2014; Yang et al. 2021). Mirror-reversal tasks are examples of *de novo* learning (Telgen et al. 2014; Yang et al. 2021). Mirror-reversal and other motor skill learning tasks are characterized by improvements in performance with practice and distinguished from adaption tasks by the presence of offline gains, i.e., performance improvements at the start of the next session relative to the end of the initial learning phase (Telgen et al. 2014; Walker et al. 2002). Motor skill learning, in general, involves changes in several brain regions, including the motor cortex (Nudo et al. 1996), basal ganglia (Yin et al., 2009), and cerebellum (Black et al. 1990). Recently, Kodama et al. (2018) showed significant increases in MRI-detected gray matter in the primary motor and somatosensory cortices and hippocampus after the first day of mirror-reversal learning, which predicted performance on later learning.

In the present study, we asked whether exposure to adaptation learning can improve subsequent *de novo* motor skill learning. There is already some evidence that prism adaptation can ameliorate symptoms of spatial neglect (Newport and Schenk 2012). As both forms of motor learning rely on and impact the function of an overlapping network of brain areas, could prism adaptation also be exploited to improve skill learning? We therefore tested whether acquisition of a novel visuomotor mapping induced by prism glasses can improve learning of a mirror-tracing task. We chose two different forms of motor learning tasks to reduce the likelihood of anterograde interference in which the learning of one visuomotor mapping disrupts learning a subsequent (opposite) mapping (Krakauer et al. 2005; Lerner et al. 2020; Miall et al. 2004). We varied prism strength from trial to trial, reducing the reliability of perceived visual feedback, which slows the rate of adaptation (Maeda et al. 2017a). This causes the brain to have to adapt for longer than a simple constant perturbation. Prism adaptation may increase motor cortex excitability, as reflected by greater motor evoked potential mean amplitude and a steeper slope of the input-output curve following transcranial magnetic stimulation (Bracco et al. 2017; Magnani et al. 2014). We predicted that adapting to prisms in one motor task would increase the excitability of neurons and their readiness to undergo structural changes, and in turn, create an optimal state for learning a subsequent task. In this sense, having to adapt to prisms would prime the brain and facilitate learning like neuromodulation techniques do (Buch et al. 2017; Orban de Xivry and Shadmehr 2014). This could provide a non-stimulation-based approach to facilitating learning and recovery of function following injury. Thus, we had one group of participants adapt to prisms that varied, trial-to-trial, in strength during a seated pointing task, followed by learning a novel mirror-reversal tracing task. We had another group perform the seated pointing task with blank lenses prior to learning the mirror-reversal tracing task. We tested both groups a second time on the tracing task the following day.

## METHODS

### Participants

Twenty-five participants (age 22.7 ± 3.0; 11 men, 14 women; all right-hand dominant) from Simon Fraser University and the surrounding area participated in this study. Participants had no known visual diseases or neurological or musculoskeletal disorders impairing movement. No participants had previously participated in a research study using prism glasses. The Office of Research Ethics at Simon Fraser University approved the study protocol, and all participants provided written informed consent before starting the experiment.

### Experimental tasks and protocol

All participants performed a precision pointing task followed by a mirror-reversal tracing task, both being completed within ∼70 minutes. The next day, participants returned to the laboratory and performed the identical mirror-reversal tracing task again. Participants used their dominant (right) hand for each task. Details are illustrated in Figure 1.

**Figure 1:**
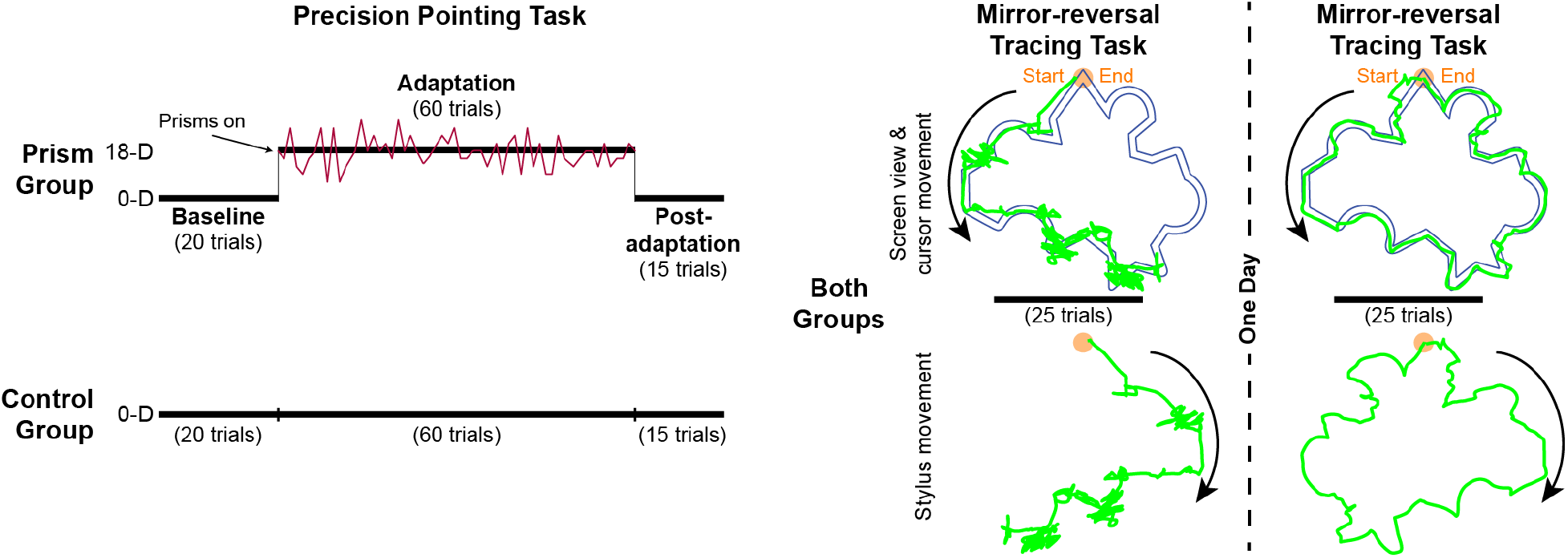
Experimental tasks and protocol. Both groups first performed 95 trials of a precision pointing task from a seated position. The control group wore goggles with 0-diopter (0-D) lenses that did not alter visual perception. The prism group wore goggles with varying strength, rightward-shifting prism lenses (mean strength = 18-diopters, 18-D) for 60 trials after pointing with 0-diopter lenses for 20 trials. This group pointed for an additional 15 trials while wearing 0-diopters lenses to test for the presence of negative aftereffects. Both groups then performed 25 trials of a tracing task in which we reversed the relationship between the stylus and cursor motion in the left-right direction, such that a rightward movement of the stylus would cause the cursor on the computer screen to move leftward. The following day, both groups repeated the tracing task. At right are examples of tracing performance (green) from one participant on trial 1 of day 1 (left) and trial 1 of day 2 (right). Top, screen view seen by the participant (colors have been changed here for clarity) with counterclockwise cursor path superimposed on the target shape (166 mm wide x 152 mm tall; path = 5 mm wide). Bottom, clockwise stylus movement. Orange circle at top of shape is tracing start- and end-point.

For the precision pointing task, participants sat with a target (vertical length = 12 cm; width = 1 cm) positioned at eye level and at a comfortable reaching distance in front. Participants kept their eyes closed except when pointing to the target. We instructed participants to reach from their chin to the target as quickly and accurately as possible, aiming for the medial-lateral (ML) center of the target. These instructions helped to reduce the possibility of online corrections of the finger during the pointing movement. We asked participants to hold their finger on the target for one second before bringing it back to their chin. An experimenter provided a verbal cue to denote the start of a trial, at which time, the participant opened their eyes to perform the movement. The vertical length of the target minimized the need for accuracy in the vertical direction because the prisms (see below) cause errors in the ML direction and our focus was ML error. An Optotrak Certus motion capture camera (Northern Digital, Waterloo, ON, Canada), placed perpendicular to the screen, recorded an infrared-emitting position marker on the medial side of the right index finger, near the tip, at a sampling frequency of 100 Hz.

We randomly assigned participants to a prism group (n = 13) or a control group (n = 12). We did not inform participants about the prism shift. Two participants in the prism group did not follow instructions, one during the pointing task and the other during the tracing task. Thus, we excluded these individuals from all analyses. All participants initially performed 20 baseline trials while wearing goggles with 0-diopter lenses. The prism group then performed 60 trials while wearing goggles with rightward-shifting prism lenses that induced a novel visuomotor mapping. We used a noise paradigm (Maeda et al. 2017a) to vary the strength of the prism lenses for each trial around a mean of 18-diopters (10.3°) and range of 6- to 30-diopters. The order of prism strengths was random except for the first and last adaptation trial, which we set to 18-diopter. The control group performed 60 trials after their baseline trials wearing 0-diopter lenses, which we removed and replaced between each trial to match the procedure for the prism group. Both groups finished with 15 pointing trials with 0-diopter lenses. For the prism group, these trials (post-adaptation phase) enabled us to test for the presence of negative aftereffects, an indication of internal model updating (Maeda et al. 2017b) and to washout any potential interference effects of the prisms on the tracing task.

After completing the precision pointing task, all participants performed 25 trials of a mirror-reversal tracing task, starting within ∼3-5 minutes after the last pointing trial. We did not inform participants about the mirror-reversal. In this task, participants sat in front of a laptop computer screen and used their dominant hand to trace between two lines forming a path around the perimeter of a shape with a stylus and tablet (Wacom Intuos Pro, medium, PTH660). We set the tablet and computer 5 cm and 31.5 cm from the edge of the table, respectively. Using Adobe Illustrator (Adobe Inc., San Jose, CA, USA), we designed an irregular shape to trace, which we imported into MATLAB (The MathWorks, Natick, MA). It comprised four semi-circles and 10 pairs of lines set at varying angles. A circle at the top of the shape marked the start and end point (see Figure 1). The width of the shape’s path measured 5 mm on the screen and 3.5 mm on the tablet. The maximum height of the shape on the screen measured 15.2 cm and the maximum width measured 16.6 cm. A custom-written MATLAB program with the Psychophysics Toolbox, version 3 (Brainard 1997) collected the tracing data. To generate the mirror reversal, we reversed the sign of the x (left-right) direction of the cursor. We instructed participants to trace the shape, moving their hand to the right first, as quickly and accurately as possible in a 90-s time frame. In addition, we instructed participants to maintain gaze on the computer screen while tracing and not to lift the stylus off the pad, as it would lose the position in the shape. We eliminated a trial if the participant moved their hand to the left initially (0.7% of trials) or if the computer did not register the stylus on the tablet (1.2% of trials). A timer in the top left corner of the screen allowed the participant and researcher to see how much time had elapsed in each trial. If they finished before 90 s, we had them keep the stylus on the start/end point and press a key to proceed to the next trial. All participants returned to the laboratory the next day to perform the same mirror-reversal tracing task again.

### Data and statistical analyses

To quantify performance on the precision pointing task, we first filtered the finger marker data using a fourth-order, low-pass Butterworth filter (cut-off frequency of 6 Hz). Next, we determined the endpoint of the pointing movement to the target, which we defined as the time at which the finger position marker’s anterior-posterior velocity profile stabilized to near zero (Bakkum et al. 2020). A total of 5.1% of trials had missing kinematic data from the finger such that we could not quantify performance. Missing data were likely the result of the participant slightly rotating their finger near the end of the pointing movement such that the position marker was no longer visible to the motion capture camera. We quantified performance (i.e., ML error) as the ML distance between the finger marker at the endpoint of the movement relative to the ML center of the target. We defined a positive value as ML error to the right, in the direction of the prism shift, and a negative value as error to the left (opposite of the shift). To determine whether participants in the prism group adapted, we compared ML error during the baseline phase (mean of the last 10 trials), first adaptation trial (or corresponding trial for the control group), late adaptation (mean of the last 10 adaptation trials or corresponding trials for the control group), and the first post-adaptation trial (or corresponding trial for the control group) using a two-way (Group x Phase) mixed-model ANOVA with participant as a random effect.

To quantify performance on the mirror-reversal task, we first used a 5-point central moving average to filter the data. We sampled the tracing data at 100 Hz. Finally, we quantified three measures: % error, completion amount, and the number of times the trace crossed the lines of the path (normalized by the completion amount; hereafter, referred to as crossing points). Specifically, we defined % error as the percentage of time the participant spent tracing outside of the path. For completion amount, we determined the centroid of the shape and converted the tracing data to polar coordinates. Subsequently, we calculated how much of the shape the participant traced, in degrees from the start position, within the 90-s time limit.

To determine how participants learned the tracing task on day 1 and whether groups learned differently, we binned the 25 trials into five five-trial averages for each of the % error, completion amount, and crossing points measures. Next, we performed separate two-way (Group x Bin) mixed-model ANOVAs (with participant as a random effect) for each measure. For the crossing points measure, we found an outlier (studentized residual of 4.7); however, removing it did not change the results and thus, we left this data point in the model. To determine the presence of offline gains with the tracing task, we compared the average of the last five trials (i.e., bin 5) on day 1 with the average of the first five trials on day 2 between groups for each of our three measures. This entailed using separate two-way (Group x Day) mixed-model ANOVAs with participant as a random effect. For this analysis, we log-transformed the % error data to conform to the assumption of normality for the ANOVA.

We used JMP 16 software (SAS Institute Inc., Cary, NC) and an alpha level of 0.05 for all statistical analyses. We conducted Tukey’s post hoc tests following a significant main effect or interaction with the ANOVAs, where appropriate. Effect sizes are reported as 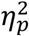.

## RESULTS

Participants in both groups performed a precision pointing task followed by a mirror-reversal tracing task. The prism group experienced rightward-shifting prism lenses of varying strengths (around a mean of 18-diopters) during a 60-trial adaptation phase of the pointing task. The control group wore 0-diopter lenses, which did not produce any perceived shift of the visual field. The next day, both groups of participants performed the identical mirror-reversal tracing task again.

### Prism adaptation

Participants pointed with their right index finger to a target positioned at eye level in front of them. Figure 2A illustrates performance across trials. The control group had an ML error of 6.3 ± 8.1 mm across repeated trials. The prism group maintained a similar performance level during baseline trials and then large ML errors in the direction of the prism shift during early adaptation. Over repeated trials, the prism group reduced errors to baseline levels. Each participant had a different prism shift order (except for the first and last adaptation trial of 18 diopters), which likely explains the variability seen in the graph across trials of the adaptation phase.

**Figure 2:**
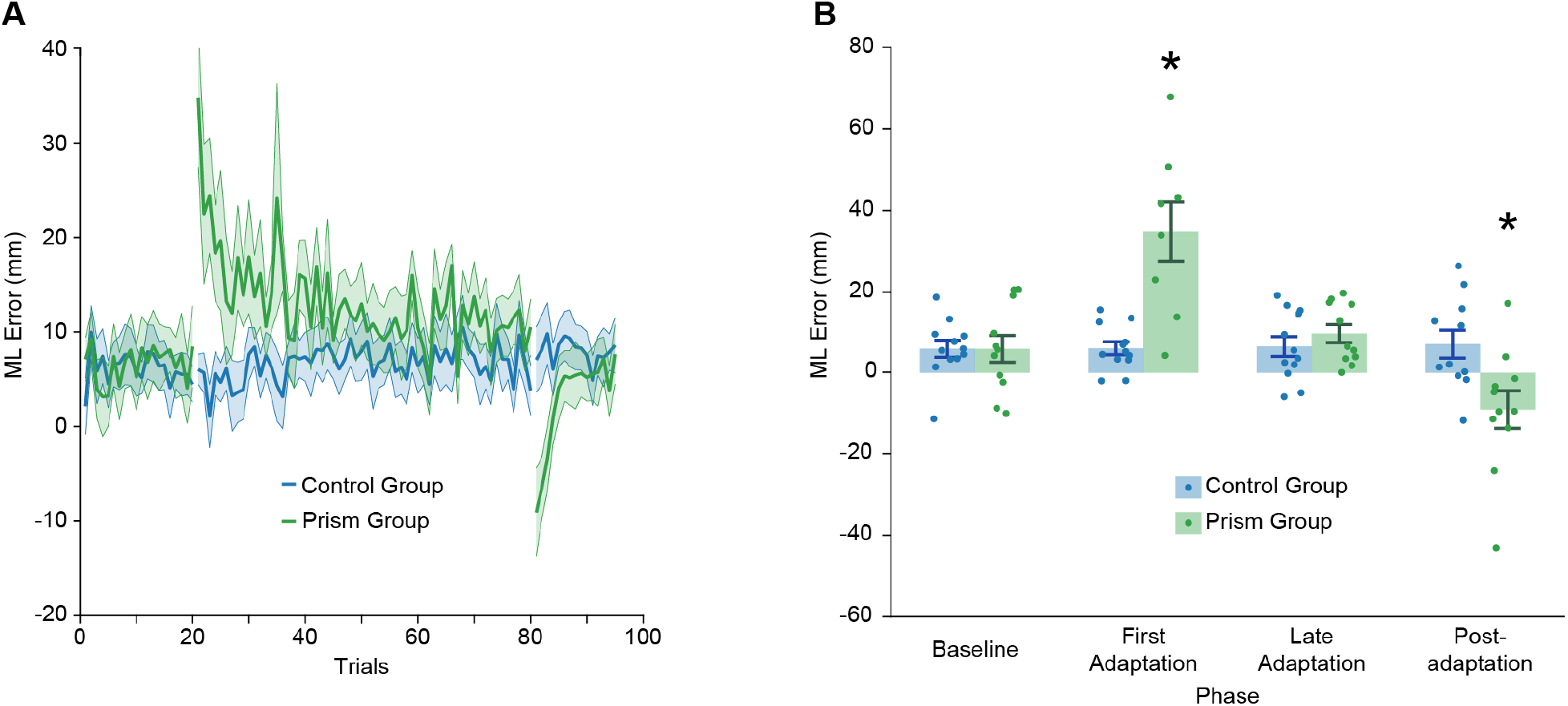
Results of the precision pointing task. **A:** Group mean ± SE, ML endpoint position error across all trials for the control (blue) and prism (green) groups. **B:** Group mean ± SE, ML endpoint position error for the baseline (or control group equivalent) phase (mean of last 10 trials), first adaptation trial (or control group equivalent), late adaptation (or control group equivalent) phase (mean of last 10 trials), and first post-adaptation trial (or control group equivalent) for the control and prism groups. For the prism group, a positive value represents errors in the direction of the prism shift. Asterisk indicates that values are significantly different from other groups/phases based on post hoc tests (p < 0.05).

To quantify group differences, we compared ML error between groups during the baseline phase (mean of the last 10 trials), first adaptation trial (or corresponding trial for the control group), late adaptation (mean of the last 10 adaptation trials or corresponding trials for the control group), and the first post-adaptation trial (or corresponding trial for the control group). We found a significant Group x Phase interaction (F_3,60_ = 21.6, p = 1.34e-9, 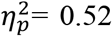). Specifically, the prism group demonstrated significantly greater error in the first adaptation phase trial compared to the control group and other phases (Figure 2B). In addition, the prism group demonstrated ML errors in the opposite direction during the first post-adaptation phase trial, indicative of a negative aftereffect. This value differed significantly from the control group and other phases.

### Mirror-reversal tracing

In the tracing task, we reversed the relationship between the stylus and cursor motion in the left-right direction, such that a rightward movement of the stylus would cause the cursor on the computer screen to move leftward. Despite this perturbation, both groups of participants demonstrated improvements in performance over repeated trials. This entailed a reduction in the time spent tracing outside the path of the shape (% error, Figure 3A), an increase in the proportion of the shape traced within the allotted time limit (completion amount, Figure 3B), and a reduction in the number of times crossing the lines of the path (normalized to completion amount; Figure 3C). Both groups maintained their performance levels on day 2, though seven control and five prism group participants required at least one trial before they could complete the shape again within the allotted time (Figure 3B).

**Figure 3:**
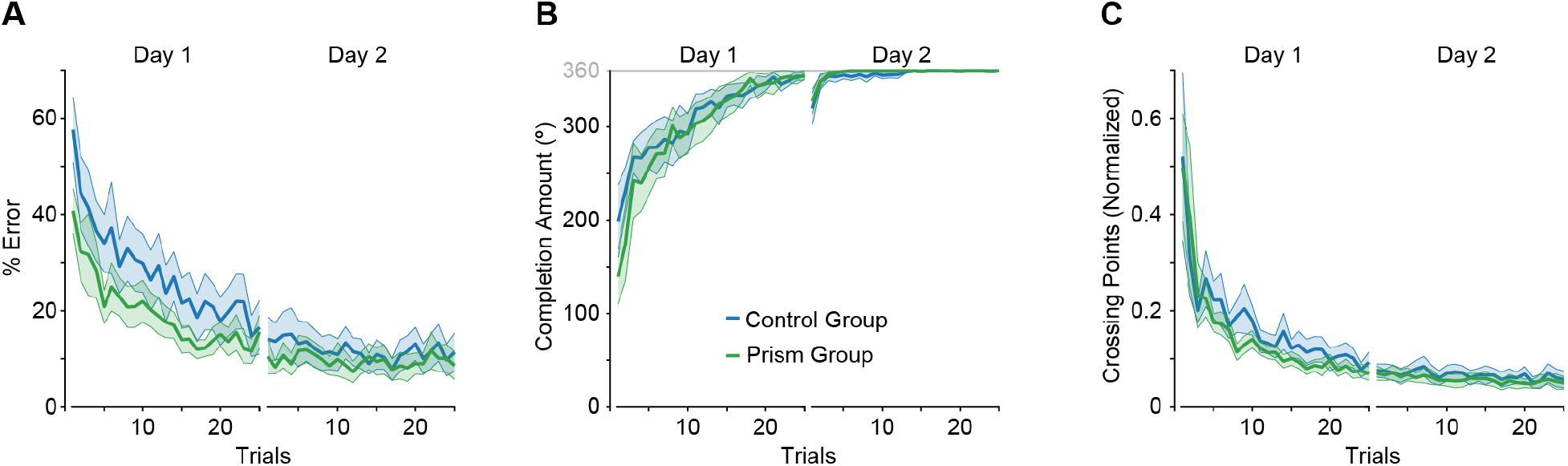
Learning profiles for the mirror-reversal tracing task. Group mean ± SE, % error (**A**), completion amount (**B**), and crossing points (**C**) across trials and days for the control (blue) and prism (green) groups.

We first asked whether groups differed in their performance on day 1. To address this question, we compared groups across five bins (each an average of five trials) for all three of our measures. For % error, we found a significant main effect of bin (F_4,84_ = 32.8, p = 1.83e-16, 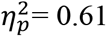), with bin 1 greater than the other bins and bin 2 greater than bins 3 – 5 (Figure 4A). However, we did not detect a significant effect of group (Group: F_1,21_ = 1.5, p = 0.241, 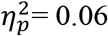; Group x Bin: F_4,84_ = 0.27, p = 0.899, 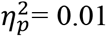). Similarly, we found a significant main effect of bin for completion amount (F_4,84_ = 38.0, p = 3.96e-18, 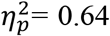), with bin 1 less than the other bins and bin 2 less than bins 3 – 5 (Figure 4B). Again, we did not detect a significant effect of group (Group: F_1,21_ = 0.18, p = 0.672, 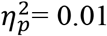; Group x Bin: F_4,84_ = 0.93, p = 0.453, 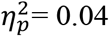). In addition, we found a significant main effect of bin for crossing points (F_4,84_ = 24.4, p = 2.02e-13, 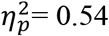), with bin 1 greater than the other bins, and bin 2 greater than bins 4 and 5 (Figure 4C). We also did not detect a significant effect of group for this measure (Group: F_1,21_ = 0.50, p = 0.487, 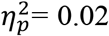; Group x Bin: F_4,84_ = 0.28, p = 0.888, 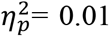). Taken together, tracing performance on day 1 showed evidence of learning but did not differ between groups.

**Figure 4:**
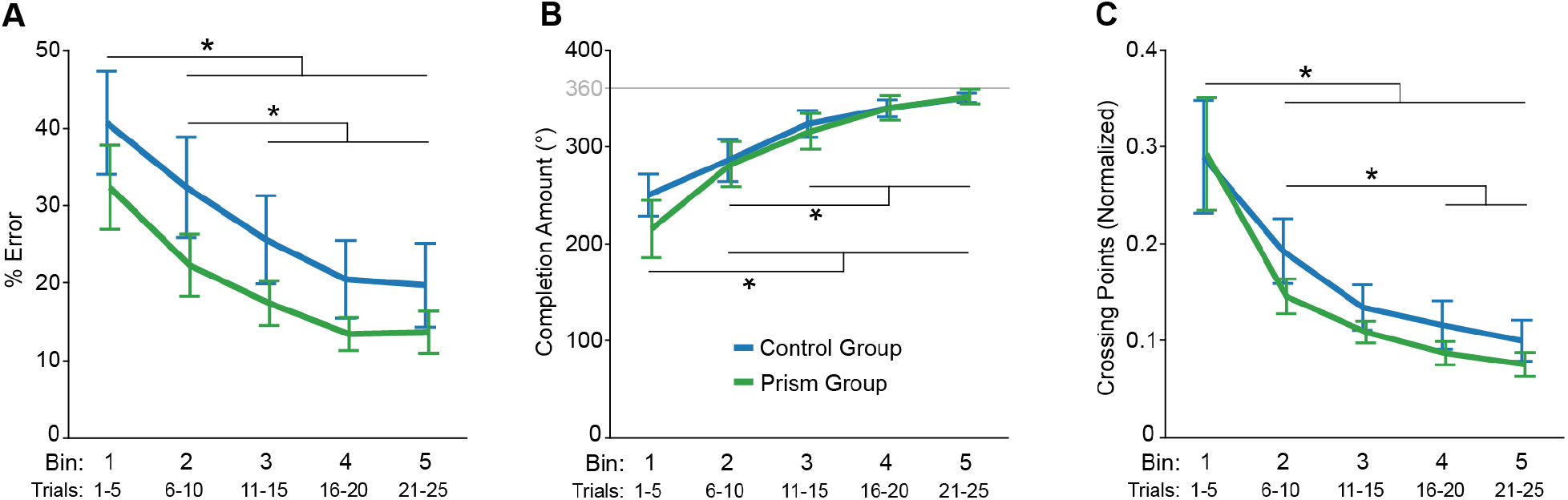
Evidence of learning on day 1 for the mirror-reversal tracing task. Group mean ± SE, % error (**A**), completion amount (**B**), and crossing points (**C**) across bins for the control (blue) and prism (green) groups. Each bin is the mean of five tracing trials for a participant, averaged within each group to produce a group mean value. Asterisk indicates that values are significantly different from each other based on post hoc tests for the main effect of bin (p < 0.05).

We next asked whether groups differed in terms of offline gains. To address this question, we compared the average of the last five trials (i.e., bin 5) on day 1 with the average of the first five trials on day 2 between groups for all three of our measures. Both groups showed reduced % error at the start of day 2 compared to the end of day 1 (main effect of day: F_1,21_ = 21.9, p = 1.27e-4, 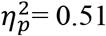; Figure 5A), supporting the notion of offline gains. However, we did not detect an effect of group (Group: F_1,21_ = 0.02, p = 0.886, 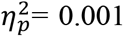; Group x Bin: F_1,21_ = 0.55, p = 0.466, 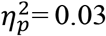). As illustrated in Figure 5B, completion amount did not differ between the start of day 2 and the end of day 1 (Day: F_1,21_ = 0.93, p = 0.345, 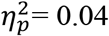) or between groups (Group: F_1,21_ = 0.1, p = 0.757, 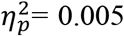; Group x Day: F_1,21_ = 0.11, p = 0.748, 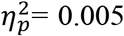). In fact, Figure 3B shows that, on average, both groups did not complete tracing the shape in the first two trials of day 2. This suggests that participants required some initial context to recall the motor memory of the visuomotor mapping and/or that they sacrificed tracing speed to maintain accuracy (note that % error was minimal on these day 2 trials; Figure 5A). Both groups showed reduced crossing points at the start of day 2 compared to the end of day 1 (main effect of day: F_1,21_ = 13.8, p = 0.001, 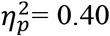; Figure 5C). We did not detect a main effect of group (F_1,21_ = 0.44, p = 0.516, 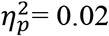) for this measure, but we found a trend towards a Group x Day interaction (F_1,21_ = 4.3, p = 0.052, 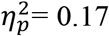). Taken together, we found evidence of offline gains but no group differences.

**Figure 5:**
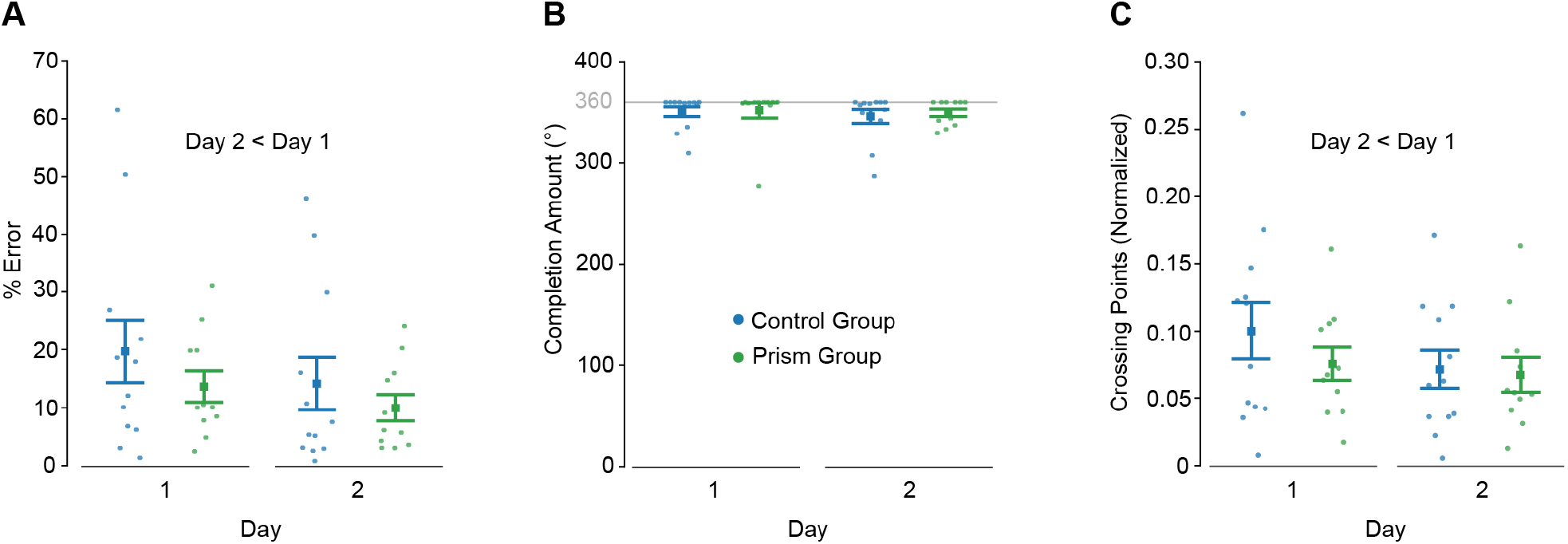
Offline gains analysis. Group mean ± SE, % error (**A**), completion amount (**B**), and crossing points (**C**) comparing the end of day 1 (mean of last five tracing trials) with the start of day 2 (mean of first five tracing trials) between groups.

## DISCUSSION

The brain undergoes extensive neuroplasticity when adapting to prisms during movement. Here, we tested whether adapting to prisms could improve the subsequent learning of a mirror-reversal tracing task. We varied the prism strength from trial to trial because it slows the rate of adaptation (Maeda et al. 2017a), thus causing the brain to have to adapt for longer than a simple constant perturbation. We predicted that adapting to these prism shifts would increase the excitability of neurons and their readiness to undergo structural changes, and in turn, create an optimal state for learning the subsequent tracing task. This is based on the premise that many of the brain regions involved in prism adaptation, including the posterior parietal cortex, cerebellum, and motor cortex, are likely also involved in learning the tracing task. Two groups of participants performed a seated pointing task, one wearing blank lenses and the other adapting to prisms. Next, they learned a novel mirror-reversal tracing task. We tested each group again the following day on the tracing task. Both groups showed similar improvements in performance on the tracing task over repeated trials on day 1. In addition, both groups showed offline gains in terms of a reduction of the percentage of time tracing outside of the shape’s path and the number of times the trace crossed the lines of the path (normalized by shape completion amount). Taken together, we did not find evidence that prior prism adaptation enhanced learning of the tracing task. Importantly, however, prior prism adaptation did not interfere with learning the tracing task.

Our tracing results showed evidence of offline gains and persistent steady-state accuracy, suggestive of a floor effect. Offline gains are characteristic of mirror-reversal tracing (Telgen et al. 2014) as well as motor sequence learning tasks (Censor et al. 2010; Fischer et al. 2002; Walker et al. 2002). We found offline gains (reduced % error and crossing points) in the tracing task in both the control and prism groups. In fact, both groups spent very little time tracing outside of the shape’s path or crossing the lines of the shape on day 2; % error and crossing point measures hovered close to zero. It is possible that the relatively small sample size of our study limited our ability to detect group differences. However, group sizes of twelve are typical in these types of studies. It is also possible that our measures suffered from somewhat of a floor effect. Nonetheless, both groups demonstrated strong evidence of learning.

Adapting to prisms during a seated pointing task prior to the mirror-reversal tracing task did not interfere with initial learning of this latter task. Some task combinations do cause anterograde interference of acquisition, especially those with similar context but conflicting or opposite mappings (e.g., Krakauer et al. 2005; Lerner et al. 2020; Miall et al. 2004). The apparent absence of interference in our experiment was likely due to differences in learning between these two tasks; visuomotor adaptation requires updating of an existing control policy whereas mirror-reversal tracing requires learning a new control policy (Krakauer et al. 2019; Sternad 2018; Telgen et al. 2014; Yang et al. 2021). Others have shown that people can learn different motor tasks concurrently. For example, learning novel kinematics (i.e., visuomotor rotations) and novel dynamics (in the form of added mass attached to the arm or reaching with a velocity-dependent force applied to the arm) do not interfere with one another (Krakauer et al. 1999; Rabe et al. 2009). Alternatively, the presence of a washout block of trials following the prism adaptation could have prevented interference (Krakauer et al. 2005).

Some mechanisms of motor learning may inhibit retention of a subsequently learned task, potentially counteracting the effect we sought to test. Long-term potentiation-(LTP)-like plasticity in human motor cortex is transiently reduced, or occluded, following simple motor learning tasks (Cantarero et al. 2013; Rosenkranz et al. 2007; Ziemann et al. 2004). This implies a finite capacity for plasticity for some time after prior learning, which could impair subsequent learning. Indeed, even the retention of the same visuomotor adaptation task is impaired on subsequent learning, although reacquisition of the task is not impacted (Hamel et al. 2020). However, our findings of high day 2 performance and offline gains in the prism group suggest that this effect did not impact retention of our tracing task. We speculate that the lack of LTP-like occlusion effect may relate to different subsets of neurons within primary motor cortex (or other learning-related brain regions) involved in our motor tasks.

There are several possible reasons why prior prism adaptation may not have enhanced subsequent *de novo* learning, which provide potential avenues for future research. First, the choice of a prism noise protocol rather than a constant prism diopter for the visuomotor perturbation may have influenced our results. Indeed, studies showing that prism adaptation improves symptoms of spatial neglect use a constant prism diopter (Newport and Schenk 2012). However, adaptation occurs more rapidly with this design compared to a noise protocol (Maeda et al. 2017a), thus reducing the time the brain must learn the mapping. It is unclear whether there are differences in metrics of plasticity between a noise and constant prism protocol. It is also possible that exposing participants to alternating blocks of trials with large prism shifts in opposite directions would have better facilitated subsequent learning. Second, we may not have used the best adaptation task (i.e., pointing with the hand) to optimize the potential of prior prism exposure. There is evidence that adapting to prisms while walking and stepping onto a target leads to stronger consolidation than reaching to a target with the arm (Bakkum et al. 2021). In addition, exposing participants to multiple tasks while adapting to prisms might have led to greater effects on the tracing task. Third, the choice of *de novo* learning task might explain the lack of group differences. Although prism adaptation did not enhance subsequent mirror-reversal tracing, it may enhance other motor skills, such as learning how to play an instrument or a certain sports skill. Ultimately, we believe our experimental rationale is sound and thus, these suggested future experiments should be conducted before concluding that prior prism adaptation (or any prior motor learning) is not beneficial to subsequent motor skill learning. Moreover, seeking such an effect is worthwhile because there are advantages to having a cheap, non-invasive, and non-stimulation-based approach to facilitate learning and recovery of function following injury.

## ACKNOWLEDGEMENTS

The authors wish to thank Julia Cielecka for help with planning and collecting pilot data, Justin Wang for programming the tracing task, and Mohammadamin Nikmanesh and Milad Hafezi for building an earlier version of the tracing task apparatus for pilot testing.

## Notes

**Disclosures:** The authors declare no conflict of interest, financial or otherwise.

### Competing Interest Statement

The authors have declared no competing interest.

## REFERENCES

Bakkum A, Donelan JM, Marigold DS. Challenging balance during sensorimotor adaptation increases generalization. J Neurophysiol. 2020; 123: 1342–1354.

Bakkum A, Donelan JM, Marigold DS. Savings in sensorimotor learning during balance-challenged walking but not reaching. J Neurophysiol. 2021; 125: 2384–2396.

Bakkum A, Marigold DS. Learning from the physical consequences of our actions improves motor memory. eNeuro. 2022; 9:eneuro.0459-21.2022.

Black JE, Isaacs KR, Anderson BJ, Alcantara AA, Greenough WT. Learning causes synaptogenesis, whereas motor activity causes angiogenesis, in cerebellar cortex of adult rats. Proc Natl Acad Sci. 1990; 87: 5568–5572.

Bracco M, Mangano GR, Turriziani P, Smirni D, Oliveri M. Combining tDCS with prismatic adaptation for non-invasive neuromodulation of the motor cortex. Neuropsychologia. 2017; 101: 30–38.

Brainard DH. The Psychophysics Toolbox. Spat Vis. 1997; 10: 433–436.

Buch ER, Santarnecchi E, Antal A, Born J, Celnik PA, Classen J, et al. Effects of tDCS on motor learning and memory formation: a consensus and critical position paper. Clin Neurophysiol. 2017; 128: 589–603.

Cantarero G, Tang B, O’Malley R, Salas R, Celnik P. Motor learning interference is proportional to occlusion of LTP-like plasticity. J Neurosci. 2013; 33: 4634–4641.

Censor N, Dimyan MA, Cohen LG. Modification of existing human motor memories is enabled by primary cortical processing during memory reactivation. Curr Biol. 2010; 20: 1545–1549.

Fischer S, Hallschmid M, Elsner AL, Born J. Sleep forms memory for fingers skills. Proc Natl Acad Sci. 2002; 99: 11987–11991.

Hamel R, Dalliare-Jean L, De La Fontaine É, Lepage JF, Bernier PM. Learning the same motor task twice impairs its retention in a time- and dose-dependent manner. Proc R Soc B. 2020; 288: 20202556.

Kodama M, Ono T, Yamashita F, Ebata H, Liu M, Kasuga S, Ushiba J. Structural gray matter changes in the hippocampus and the primary motor cortex on an-hour-to-one-day scale can predict arm-reaching performance improvement. Front Hum Neurosci. 2018; 12:209.

Krakauer JW, Ghez C, Ghilardi MF. Adaptation to visuomotor transformations: consolidation, interference, and forgetting. J Neurosci. 2005; 25: 473–478.

Krakauer JW, Ghilardi M-F, Ghez C. Independent learning of internal models for kinematic and dynamic control of reaching. Nat Neurosci. 1999; 2: 1026–1031.

Krakauer JW, Hadjiosif AM, Xu J, Wong AL, Haith AM. Motor learning. Compr Physiol. 2019; 9: 613–663.

Landi SM, Baguear F, Della-Maggiore V. One week of motor adaptation induces structural changes in primary motor cortex that predict long-term memory one year later. J Neurosci. 2011; 31: 11808–11813.

Lerner G, Albert S, Caffaro PA, Villalta JI, Jacobacci F, Shadmehr R, Della-Maggiore V. The origins of anterograde interference in visuomotor adaptation. Cer Cortex. 2020; 30: 4000–4010.

Maeda RS, O’Connor SM, Donelan JM, Marigold DS. Foot placement relies on state estimation during visually guided walking. J Neurophysiol. 2017a; 117: 480–491.

Maeda RS, McGee SE, Marigold DS. Consolidation of visuo-motor adaptation memory with consistent and noisy environments. J Neurophysiol. 2017b; 117: 316–326.

Maeda RS, McGee SE, Marigold DS Long-term retention and reconsolidation of a visuomotor memory. Neurobiol Learn Mem. 2018; 155: 313–321.

Magnani B, Caltagirone C, Oliveri M. Prismatic adaptation as a novel tool to directionally modulate motor cortex excitability: evidence from paired-pulse TMS. Brain Stim. 2014; 7: 573–579.

McDougle SD, Ivry RB, Taylor JA. Taking aim at the cognitive side of learning in sensorimotor adaptation tasks. Trends Cogn Sci. 2016; 20: 535–544.

Miall RC, Jenkinson N, Kulkarni K. Adaptation to rotated visual feedback: a re-examination of motor interference. Exp Brain Res. 2004; 154: 201–210.

Newport R, Schenk T. Prisms and neglect: what have we learned? Neuropsychologia. 2012; 50: 1080–1091.

Nudo RJ, Miliken GW, Jenkins WM, Merzenich MM. Use-dependent alterations of movement representations in primary motor cortex of adult squirrel monkeys. J Neurosci. 1996; 16: 785–807.

O’Shea J, Revol P, Cousijn H, Near J, Petitet P, Jacquin-Courtois S, Johansen-Berg H, Rode G, Rossetti Y. Induced sensorimotor cortex plasticity remediates chronic treatment-resistant visual neglect. eLife. 2017; 6:e26602.

Orban de Xivry J-J, Shadmehr R. Electrifying the motor engram: effects of tDCS on motor learning and control. Exp Brain Res. 2014; 232: 3379–3395.

Panico F, Rossetti Y, Trojano L. On the mechanisms underlying prism adaptation: a review of neuro-imaging and neuro-stimulation studies. Cortex. 2020; 123: 57–71.

Panico F, Sagliano L, Sorbino G, Trojano L. Engagement of a parietal-cerebellar network in prism adaptation. A double-blind high-definition transcranial direct current stimulation study on healthy individuals. Cortex. 2022; 146: 39–49.

Petitet P, O’Reilly JX, O’Shea J. Towards a neuro-computational account of prism adaptation. Neuropsychologia. 2018; 115: 188–203.

Rabe K, Livne O, Gizewski ER, Aurich V, Beck A, Timmann D, Donchin O. Adaptation to visuomotor rotation and force field perturbation is correlated to different areas in patients with cerebellar degeneration. J Neurophysiol. 2009; 101: 1961–1971.

Redding GM, Rossetti Y, Wallace B. Applications of prism adaptation: a tutorial in theory and method. Neurosci Biobehav Rev. 2005; 29: 431–444.

Rosenkranz K, Kacar A, Rothwell JC. Differential modulation of motor cortical plasticity and excitability in early and late phases of human motor learning. J Neurosci. 2007; 27: 12058–12066.

Schintu S, Freedberg M, Gotts SJ, Cunningham CA, Alam ZM, Shomstein S, Wassermann EM. Prism adaptation modulates connectivity of the intraparietal sulcus with multiple brain networks. Cerebral Cortex. 2020; 30: 4747–4758.

Sternad D. It’s not (only) the mean that matters: variability, noise and exploration in skill learning. Curr Opin Behav Sci. 2018; 20: 183–195.

Telgen S, Parvin D, Diedrichsen J. Mirror reversal and visual rotation are learned and consolidated via separate mechanisms: recalibrating or learning De Novo? J Neurosci. 2014; 34: 13768–13779.

Tseng YW, Diedrichsen J, Krakauer JW, Shadmehr R, Bastian AJ. Sensory prediction errors drive cerebellum-dependent adaptations of reaching. J Neurophysiol. 2007; 98: 54–62.

Walker MP, Brakefield T, Morgan A, Hobson JA, Stickgold R. Practice with sleep makes perfect: sleep-dependent motor skill learning. Neuron. 2002; 35: 205–211.

Yang CS, Cowan NJ, Haith AM. De novo learning versus adaptation of continuous control in a manual tracking task. eLife. 2021; 10: e62578.

Yin HH, Mulcare SP, Hilário MRF, Clouse E, Holloway T, Davis MI, Hansson AC, Lovinger DM, Costa RM. Dynamic reorganization of striatal circuits during the acquisition and consolidation of a skill. Nature Neurosci. 2009; 12 333–341.

Ziemann U, Iliać T, Pauli C, Meintzschel F, Ruge D. Learning modifies subsequent induction of long-term potentiation-like and long-term depression-like plasticity in human motor cortex. J Neurosci. 2004; 24: 1666–1672.

